# Informing agent-based models with spatial data using convolutional autoencoders

**DOI:** 10.64898/2026.03.13.711699

**Authors:** B. Wang, C. Liao, E.H.J. Danen, E. Neubert, F. Eduati

**Author notes:** corresponding author <u>Contact information</u> Lead contact: Eduati F.

## Abstract

Spatial computational models such as agent-based models (ABMs) offer powerful *in silico* tools to study tumor dynamics, yet imaging data are still rarely used to inform these models directly. We present an ABM optimization framework that leverages convolutional encoders to compare spatial patterns between experimental imaging data and ABM-generated outputs within a shared latent space. This quantitative comparison was used to estimate ABM parameters across three datasets, ranging from synthetic data to 3D tumoroid–T cell co-culture microscopy and histopathology images from The Cancer Genome Atlas skin cutaneous melanoma samples. Estimated parameters were evaluated using data-derived features and experimental knowledge, including experimental conditions and gene expressions. Simulations using optimized parameters reproduced key spatial features of the training images, such as tumor boundary complexity and tumor–tumor neighborhood structure. Together, these results demonstrate a flexible framework for ABM parameter optimization using spatial data across modalities, enabling systematic investigation of how spatial architecture influences tumor progression and immune interactions.

## Introduction

The tumor microenvironment (TME) is a complex biological system, in which spatial organization plays a critical role in shaping tumor development and therapeutic response (Lindsay et al., 2025; Eliason et al., 2025; Fassler *et al*., 2022). Cellular location and interactions, particularly at the tumor–immune interface, drive tumor behavior and immune evasion, and spatial features have shown promise as biomarkers for cancer progression, therapy response, and recurrence (Lindsay *et al*., 2025; Eliason et al., 2025; Fassler et al., 2022).

Computational agent-based models (ABMs) provide a mechanistic framework to study how spatial organization emerges from cell behaviors and interactions, and how it influences tumor progression and therapeutic response (West et al., 2023; Stephan *et al*., 2024). ABMs define rules governing the behavior of individual agents (e.g., proliferation, death) and their interactions (e.g., migration, immune cell–mediated tumor cell killing). Based on these rules, ABMs stochastically simulate the spatial evolution of tumors, capturing how local interactions give rise to complex spatial organization over time. These rules are governed by parameters that quantify the rates and strengths of cellular behaviors and interactions.

However, a key limitation of ABMs lies in parameterization. As ABM parameters are biologically interpretable by construction, some can be fixed to plausible values based on literature or directly measured experimentally. Some parameters however might be unknown or highly context specific and require estimation from experimental data, most commonly using particle swarm optimization (PSO) (Cogno et al., 2024). These approaches rely on fitting model outputs to quantitative experimental measurements. For example, migration rates have been directly informed by scratch assays (Kather *et al*., 2017), whereas other parameters have been estimated by fitting model simulations either to time resolved measurements of tumor cell abundance under different treatment conditions, such as Incucyte data (van Genderen et al., 2024), or to steady state histology derived cell types ratios (Kather *et al*., 2018).

However, none of these data types explicitly capture the rich spatial information contained in tissue images that could be used to inform ABMs. While recent advances in spatial imaging technologies and AI based analysis of standard pathology images enable increasingly detailed quantification of cellular composition and organization within tumors (Booij et al., 2019; Rios and Clevers, 2018; Lapuente-Santana et al., 2024), leveraging such data for quantitative comparison with ABM simulations remains challenging. This difficulty arises from the high dimensionality of imaging data, the diversity of imaging modalities, and the complex mapping between images and computational simulations. One approach has been to extract predefined spatial features from images for model calibration (Macklin et al., 2012), but this captures only a subset of the available spatial information and imposes a priori assumptions on which spatial patterns are considered relevant. Representation learning offers an alternative by deriving low-dimensional embeddings of experimental images, into which simulated ABM outputs can be projected for quantitative comparison during parameter optimization. Neural networks such as autoencoders are particularly effective in this context, capturing complex nonlinear patterns beyond classical approaches like principal component analysis (Wani, 2025).

Cess and Finley pioneered the use of representation learning for ABM model calibration, demonstrating that learned embeddings can bridge fluorescence tumor imaging data and simulations (Cess and Finley, 2023). However, the generalizability of this strategy across imaging modalities and its implications for parameter identifiability and biological interpretation have not been systematically investigated. Here, we develop an adaptable autoencoder-based framework that addresses these challenges and evaluates parameter estimation across synthetic, in vitro, and clinical contexts. Using a simple architecture with two convolutional layers, we demonstrate robust image comparison between simulations and experimental data across three datasets: synthetic ABM configurations, 3D tumor–T cell coculture microscopy, and TCGA histopathology. Together, these results provide a systematic assessment of embedding-based calibration and establish a foundation for linking image-derived spatial features to mechanistic model parameters.

## Materials and methods

### Agent-based model (ABM)

We implemented a 2D grid-based ABM comprising tumor cells and lymphocytes as agents, adapting model rules and parameter definition from (Kather *et al*., 2018). Each grid position can be occupied by a single agent. Agents act asynchronously in random order at each simulation step according to biologically interpretable rules governing proliferation, death, migration, lymphocyte influx, and immune-mediated tumor cell killing. These rules are controlled by parameters that quantify the rates and probabilities of cellular behaviors and interactions. Unless otherwise specified, all parameters were fixed to the values reported in the **Supplementary Table**. We focused on several parameters expected to be context specific and therefore estimated from data: tumor proliferation (TUpprol), lymphocyte-mediated tumor killing probability (IMpkill), and parameters governing lymphocyte access to the tumor, including random walk probability (IMrwalk), which controls directed versus random migration, and lymphocyte influx probability (IMinfluxProb), which regulates immune cell recruitment into the system.

### Data sets and image-to-ABM mapping

All datasets were processed to obtain spatial maps of tumor and lymphocyte distributions compatible with the ABM lattice representation.

#### Synthetic simulations

We generated 3,000 parameter settings in which TUpprol and IMpkill were independently sampled with Latin hypercube sampling over [0.1, 0.5] and [0.1, 0.8], respectively, and combined with three discrete IMrwalk values (0, 0.5, 1), corresponding to fully directed to fully random lymphocyte migration. For each parameter setting, 10 stochastic replicates were simulated, resulting in a total of 30,000 simulations. Simulations were performed on a 100×100 grid, initialized from the same configuration consisting of 116 tumor cells and 6 lymphocytes. Each simulation was run for 150 steps, and only the final simulation states were used for downstream analyses.

#### Co-culture microscopy images

3D tumoroid–T cell co-culture images were obtained from Liao et al. using timelapse confocal microscopy acquired every 1 hour over 72 hours. The data set contains one time-lapse per Z stack, acquired every 10 μm (Liao *et al*., 2025). Two experimental conditions were included, corresponding to experiments with effective bispecific antibodies (bsAb, M07) or non-effective bsAb (M10), for a total of 1,152 images from 16 timelapses. Images were downsampled to 100×100 resolution and directly mapped onto the ABM grid by assigning each pixel to the cell type with the highest signal intensity above a threshold of 50. Hollow regions were filled assuming solid tumor masses using Canny edge detection (thresholds 200 and 30) followed by morphological operations (kernel size 35×35). The final image in each timelapse was used for ABM optimization.

#### Histopathology slides

TCGA skin cutaneous melanoma slides (n=366) with cell probability maps generated by SPoTLighT (Lapuente-Santana et al., 2024) were filtered for desert and immune-enriched non-fibrotic subtypes (Bagaev *et al*., 2021). Images were divided into four patches, selecting those with at least ≥80% coverage, resulting in a total of 292 TCGA patches (162 immune desert and 130 immune-enriched). Patches were mapped onto 25×25 ABM grids, and cell types (tumor or lymphocyte) were assigned for probabilities >0.5. Corresponding bulk RNA-seq data (https://gdac.broadinstitute.org) were retrieved for comparison with optimized ABM parameter values.

### Autoencoder

An autoencoder was employed to learn a low-dimensional embedding of input images, enabling spatial patterns in experimental data and ABM simulations to be compared in a common latent space for parameter estimation. The encoder consisted of two 3×3 convolutional layers with 2×2 max pooling and ReLU activations, followed by flattening to a latent vector of size 256. The decoder mirrored the encoder architecture and ended with a sigmoid activation. The network was trained using the Adam optimizer with mean squared error loss and batch sizes of 32 (TCGA), 64 (tumoroid), and 128 (synthetic) for 3000 iterations. Because reconstructed images from the autoencoder exhibited smoothing relative to the original input values, blurring the distinction of cell types, we discretized the output images into three values representing tumor cells, lymphocytes, and unoccupied grid cells. This discretization enabled direct comparison between reconstructed images and training samples.

### ABM optimization

ABM parameters were optimized by minimizing mean squared error between latent vectors of simulations and experimental data using particle swarm optimization (PSO; 12 particles, 3 iterations, c1=2, c2=1, w=0.5) (Stephan et al., 2024). For our analyses, we use the best estimate defined by a lowest error value from the PSO. The number of stochastic replicates per dataset was selected to balance variation reduction and computational efficiency (synthetic=2, tumoroid=20, TCGA=5). See **Supplementary Table** for fixed parameter values used for each data set. Initial configurations matched the initial spatial mapping from the synthetic data (synthetic, TCGA) or initial microscopy timelapse image (tumoroid).

### ABM validation

We compared estimated parameter values with properties from training data (i.e. original parameter values for synthetic data) with Pearson correlations (α=0.05) and Wilcoxon rank sum tests (for pairwise comparisons, i.e. experimental conditions for tumoroid images, gene expression for TCGA patches) with Benjamini-Hochberg correction. We also analyzed spatial features through two spatial metrics:

1. **Complexity score (equation 1)**: normalized measure of spheroid roughness (Beslmüller et al., 2025; Hou et al., 2018) A circle has a complexity of 1, and irregular shapes will translate to a higher complexity measure. We compared complexity scores only for clearly defined spheroids (tumoroid, synthetic with TUpprol < 0.3) and further filtered calculated values based on how well edges were detected for the perimeter and area. TCGA patches and synthetic data with TUpprol > 0.3 were excluded, as the image often consisted of a densely filled grid for which calculated perimeter and area would be distorted due to the main tumor mass having reached the grid boundaries.
2. **Tumor neighbor count**: mean number of tumor and lymphocyte neighbors per tumor cell, based on a graph representation of the ABM spatial output (Lapuente-Santana *et al*., 2024). The edges of the graph were assigned based on a Moore neighborhood (radius 1).

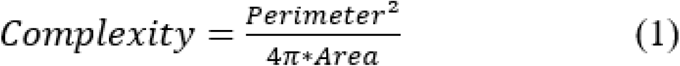

## Results

### Training convolutional autoencoders on different imaging data

We prepared three datasets mapped onto the ABM grid (**Methods, Figure 1A**), comprising 30,000 synthetic images, 1,152 tumoroid microscopy images from two experimental conditions, and 292 TCGA histopathology patches representing two immune subtypes. Separate convolutional autoencoders were trained on each dataset using a 90/10 train–validation split. Complexity scores were moderately correlated between original and reconstructed images (synthetic: R = 0.41, p<2.2e-16; tumoroid: R = 0.27, p=0.0037; **Figure 1B**), with especially tumoroid reconstructions showing an overall reduction in complexity consistent with smoothing. Tumor–tumor neighbor relationships were well preserved (R = 0.80, 0.93, 0.92; p<6e-13; **Figure 1C**), in contrast to differing correlations found for tumor–lymphocyte neighbors. While tumor-lymphocyte neighbors were highly preserved in TCGA patches (R = 0.96, p < 2.2e-16) and still captured moderately in tumoroid data (R = 0.54, p = 6.1e-10), a negative correlation was present for synthetic data (R = -0.34, p < 2.2e-16). The best alignment in tumor-lymphocyte neighbors for TCGA could reflect the presence of more contiguous lymphocyte regions, whereas lymphocytes appear more sparsely distributed in tumoroid and synthetic images and are therefore harder to preserve during reconstruction (**Supplementary Figure 1**).

**Figure 1.**
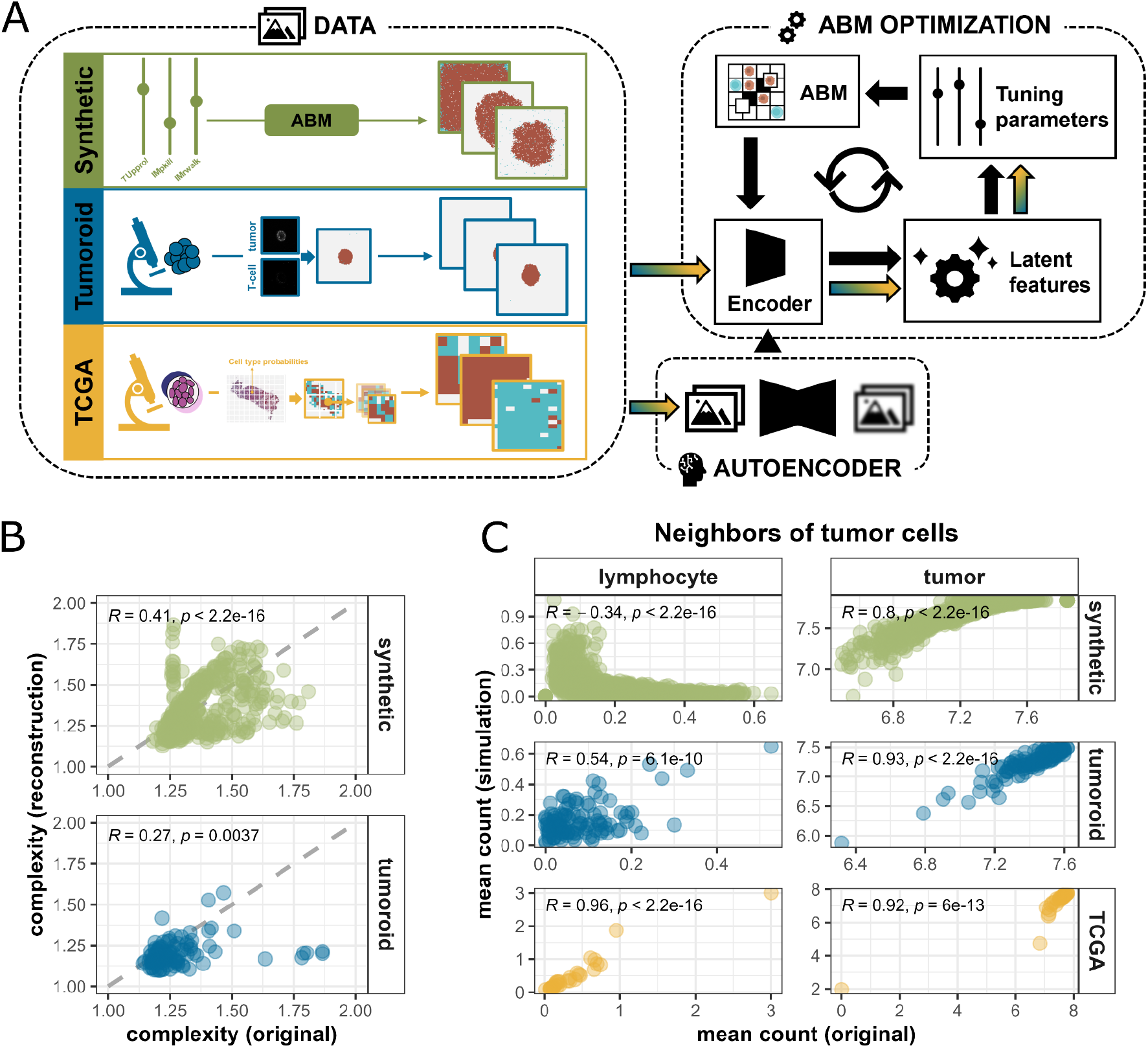
Pipeline overview and autoencoder. **(A)**. Schematic of the pipeline for ABM parameter estimation, showing three datasets and preprocessing steps. **(B)** Comparison of complexity scores between original and autoencoder-reconstructed images in the validation set; Pearson correlations shown, dashed line indicates x=y. **(C)** Comparison of tumor cell neighbor counts between original and reconstructed images; Pearson correlations shown for the validation set.

Next, we estimated ABM parameters for 75 synthetic images, 16 tumoroid timelapses (different Z stacks per experiment, training image is the final image per timelapse) and 292 TCGA patches, using the encoder layers from trained autoencoders (**Figure 1**). Unless otherwise stated, the autoencoders were trained separately for each data modality and then applied to the corresponding experimental data and model simulations for optimization of model parameters. Model performance was evaluated by comparing estimated parameters with dataset-derived information and spatial similarity between original and simulated images.

### Synthetic data enables systematic investigation of ABM optimization results

For synthetic data, estimated parameters were compared with ground-truth values, optimizing tumor proliferation (TUpprol), lymphocyte-mediated tumor killing (IMpkill), and lymphocyte random walk (IMrwalk) (**Figure 2A**). TUpprol was accurately recovered (R = 0.92, p < 2.2e-16), whereas IMpkill was poorly estimated (R = 0.06, p = 0.61). IMrwalk showed small, non-significant differences across values, suggesting that TUpprol alone largely determines spatial patterns in the simulations.

**Figure 2.**
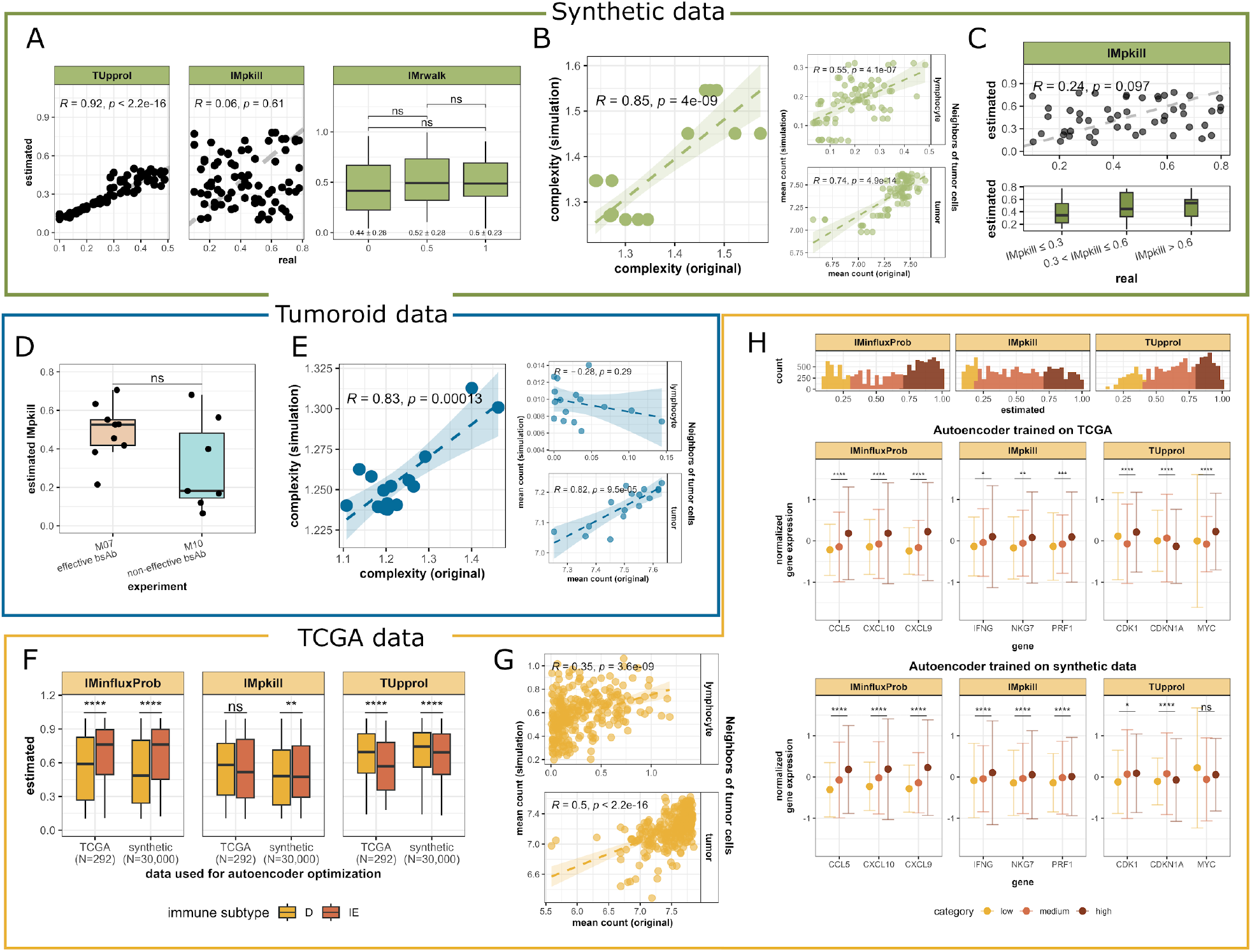
ABM parameter optimization with different imaging data sets. **(A)** Comparison of estimated vs. ground-truth ABM parameters using synthetic data: tumor proliferation (TUpprol), lymphocyte-mediated killing (IMpkill), and lymphocyte random walk (IMrwalk). Pearson correlations are shown for TUpprol and IMpkill; Wilcoxon tests with Benjamini-Hochberg correction for IMrwalk, including means ± SD. **(B)** Comparison of complexity scores and spatial fidelity (tumor-tumor or tumor-lymphocyte neighbors) of original vs. simulated images for synthetic data. **(C)** Independent IMpkill estimation with fixed TUpprol and IMrwalk. A comparison between ground-truth and estimated IMpkill is given for both continuous values (top) and discretized values (bottom). **(D)** Estimated IMpkill values of tumoroid images across experimental conditions. **(E)** Comparison of complexity scores and spatial fidelity (tumor-tumor or tumor-lymphocyte neighbors) of original vs. simulated images for tumoroid data. **(F)** ABM parameters estimated with TCGA data (IMinfluxProb (lymphocyte influx), IMpkill, TUpprol) per immune subtype (desert vs. immune-enriched) and optimized using autoencoders trained on different data sets (TCGA, synthetic). The means ± SD and Wilcoxon tests shown. Sizes of each data set used for the autoencoders is also provided. **(G)** Comparison of spatial fidelity (tumor-tumor or tumor-lymphocyte neighbors) of original vs. simulated images for TCGA patches. **(H)** Assessing estimated ABM parameter values for TCGA data with corresponding bulk transcriptomics. We stratified estimated parameter values into three categories (top figure) for this comparison. For each ABM parameter, we illustrate three markers involved in lymphocyte recruitment, cytotoxicity and tumor proliferation, with corresponding Wilcoxon tests (bottom figures).

We next assessed how well simulations using the optimised parameters captured spatial features of the original data. Complexity scores were strongly correlated between original and simulated images (R = 0.85, p = 4e-09; **Figure 2B**). Spatial neighbor relationships were also significantly retained (p < 4.1e-07; **Figure 1B**), with tumor–tumor neighbors (R=0.74) better captured than tumor-lymphocyte neighbors (R=0.55), consistent with the weaker recovery of lymphocyte-related parameters.

To further investigate the estimation of IMpkill within the current latent space, we reduced variability from other parameters. We fixed TUpprol to 0.1, 0.2, 0.3, 0.4, or 0.5 by selecting ten images per value (maximum deviation of 0.01) and generated with an IMrwalk of 1 to ensure directed lymphocyte migration toward tumor cells, thereby increasing tumor–lymphocyte interactions necessary to observe killing effects. Under these constraints, the correlation between estimated and ground-truth IMpkill of 50 samples increased but remained moderate and did not reach statistical significance (R = 0.24, p = 0.097; **Figure 2C**). Nevertheless, stratifying ground-truth IMpkill into three categories (≤ 0.3, > 0.3 and ≤ 0.6, and > 0.6) revealed a monotonic increase in median estimated IMpkill (0.342, 0.448, and 0.539, respectively) indicating that the latent representation allows to capture biologically meaningful trends in killing probability, despite imperfect overall recovery. Further analysis showed that discrepancies between estimated and true IMpkill varied across TUpprol values (**Supplementary Figure 2**): lower TUpprol values were associated with more spherical tumor structures, whereas higher TUpprol values resulted in densely populated grid-like configurations. The latter configuration exhibits reduced spatial variability across replicates, likely due to the smaller number of empty grid cells available to introduce stochastic variation between simulations. This distinction may partly explain the smaller variation of differences between estimated and ground-truth IMpkill observed at higher TUpprol levels.

### Differences in estimated IMpkill values reflect the experimental conditions of the tumoroid images

For the in vitro tumoroid co-culture case study, we fixed TUpprol to its empirical value because synthetic experiments indicated that TUpprol can dominate multi-parameter optimization and because both experimental conditions used the same tumor cell line, implying that proliferation rates should remain constant. e therefore estimated only IMpkill, the parameter expected to differ between conditions due to the varying effectiveness of the bispecific antibody, while keeping all other parameters fixed (see **Supplementary Table**). Moreover, as we observed from the synthetic data a larger variability in IMpkill optimization with spherical structures, we increased the number of replicates for each optimization step to 20.

IMpkill estimates were higher for M07 than for M10 (median 0.526 vs 0.182; **Figure 2D**), consistent with the expected differences between effective or non-effective bsAb conditions. Although this difference did not reach statistical significance, the limited number of samples per condition (9 for M07, 7 for M10) may have reduced statistical power. When comparing model simulations using optimized parameters with the corresponding images used for training, complexity scores were highly correlated (R = 0.83, p = 1.3e-04; F**igure 2E**), and tumor–tumor neighbor relationships were also well captured (R = 0.82, p = 9.5e-05). In contrast, tumor-lymphocyte neighbor relationships showed a weak negative correlation (R = -0.29, p=0.29). Tumoroid images exhibited the sparsest lymphocyte distributions, which, combined with latent-space compression and stochasticity in lymphocyte behavior, may have reduced the ability to reproduce these patterns in simulations. This added effect of stochasticity could also be an explanation how reconstructing images with the autoencoder still resulted in moderate results (**Figure 1B**), while these local structures were more difficult to capture from the stochastic ABM simulations.

### Optimized ABM values can be correlated with TCGA gene expression data and immune subtypes

Patches derived from TCGA pathology images represent larger in vivo tumor regions and, unlike the synthetic and tumoroid datasets, the grid is typically densely populated with tumor and lymphocyte cells, with few empty regions. Immune cells appear in spatially coherent areas rather than as isolated single cells,such that the data primarily capture differences in immune infiltration and cluster distribution. We included immune desert and immune-enriched non-fibrotic subtypes with distinct spatial patterns (**Supplementary Figure 3A-B**). To reflect differences in immune recruitment at this scale, we therefore estimated lymphocyte influx probability (IMinfluxProb) in addition to TUpprol and IMpkill.

Examination of the estimated model parameters revealed significant differences in TUpprol and IMinfluxProb between immune subtypes, with desert tumors exhibiting higher TUpprol and lower IMinfluxProb (**Figure 2F**) consistent with their biological characteristics. In contrast, IMpkill showed no subtype-specific differences. Spatial organization was also well preserved between pathology-derived cell maps and the corresponding optimized simulations, as reflected by moderate correlations in tumor–tumor and tumor–lymphocyte neighbor counts (R=5 and R=0.35, respectively; p-val < 3.6e-09; **Figure 2G**). Together, these results suggest that our framework enables personalization of ABM based on pathology imaging data. Nonetheless, the absence of a significant subtype difference in IMpkill was unexpected.

One possible explanation was that the autoencoder trained on the limited TCGA dataset (292 TCGA patches, in contrast to 30,000 synthetic or 1,152 tumoroid images), may not have learned sufficiently informative representations for robust parameter optimization. To assess this, we re-estimated TUpprol, IMpkill and IMinfluxProb using the autoencoder previously trained on synthetic data. The resulting parameter estimates again showed significant differences in IMinfluxProb and TUpprol across immune subtypes (**Figure 2F**). Interestingly, IMpkill was now significantly different between subtypes, with higher mean values in the immune-enriched subtype (0.651) compared to the desert subtype (0.546). These results suggest that an autoencoder trained purely on synthetic data can generalize to clinical pathology images and may enhance the robustness of parameter estimation when experimental data are limited.

To biologically validate the parameters estimated from pathology images, we leveraged matched bulk gene expression data to assess whether TUpprol, IMpkill, and IMinfluxProb were associated with expression of biologically related genes, including proliferation markers (CDK1, CDKN1A, MYC (Wang et al., 2023; Shen *et al*., 2025; Duffy *et al*., 2025)), cytotoxic effector genes (IFNG, NKG7, PRF1 (Wen *et al*., 2022; Guan *et* al., 2024)), and chemokines involved in lymphocyte recruitment (CCL5, CXCL10, CXCL9 (Yu *et al*., 2020; Kraja *et al*., 2025)). Samples were stratified into low, medium, and high parameter groups using thresholds derived from the estimated parameter distributions (TUpprol < 0.4 vs > 0.8; IMpkill < 0.2 vs > 0.7; IMinfluxProb < 0.3 vs > 0.7; **Figure 2H**).

Using parameter estimates obtained from both TCGA-trained and synthetic-trained autoencoders, we tested for differences in normalized gene expression between low and high parameter groups. Significant associations were observed for multiple gene–parameter pairs (**Figure 2H**). For the TCGA-trained model, higher IMinfluxProb was associated with increased expression of recruitment-related chemokines, and high IMpkill values corresponded to elevated cytotoxic gene expression compared to low IMpkill samples. Proliferation genes (CDK1, MYC) showed higher expression in high-TUpprol samples, while the proliferation suppressor CDKN1A exhibited the expected inverse association.

Similar trends were observed using the synthetic-trained autoencoder. Notably, subtype differences in IMpkill were more pronounced, whereas associations between TUpprol and proliferation-related genes were weaker for CDK1 and did not reach statistical significance for MYC (**Figure 2H**). The similarity for IMinfluxProb and TUpprol, while IMpkill resulted in more distinct results, is also reflected in how well estimated parameter values correlated between ABM parameters estimated using either the autoencoder trained on TCGA or synthetic data (**Supplementary Figure 3C**). Overall, these results indicate that ABM parameters inferred from standard pathology images capture biologically meaningful processes, demonstrating that embedding-based calibration enables quantitative interpretation of tumor proliferation and immune activity from spatial imaging data.

## Discussion

We developed an agent-based model (ABM) optimization pipeline in which parameters are informed directly by spatial data. By using convolutional encoders to compress spatial features into a low-dimensional latent space, the framework can be applied across diverse imaging data modalities, from purely *in silico* studies to confocal microscopy or histopathology images. When images are mapped onto the ABM grid with tumor and lymphocyte placements, the pipeline enables estimation of ABM parameters that capture both data-derived characteristics and spatial architecture, such as spheroid roughness and spatial fidelity. We further demonstrate that these parameters align with biologically meaningful features derived from independent molecular data, reinforcing the mechanistic interpretability of the framework. Finally, our results suggest that synthetic data can support parameter optimization in data-limited settings, highlighting its potential to enhance robustness and generalizability of spatial model calibration.

While varying accuracy was achieved in estimating parameter values of the synthetic data set, local features such as tumor-tumor or tumor-lymphocyte neighbors were still adequately represented. These results suggest that different combinations of parameters could yield similar spatial output. Parameter identifiability and correlation analyses would therefore be important in defining the set of parameters to optimize and their corresponding boundary values. These analyses are however not trivial. An important reason is the inherent stochasticity of ABMs, making them well suited for modeling tumor–immune interactions but posing challenges for parameter optimization and identifiability (Cess and Finley, 2023; Hamis *et al*., 2020). Therefore, increasing the number of simulations per parameter set, more extensive, downstream analyses and systematic hyperparameter optimization would be essential to improve parameter estimation and derive deeper biological insights. Another interesting input that could be varied is the initial configuration and characterizing how this impacts the resulting spatial outputs.

A promising future direction is to test alternative representation learning approaches for mapping spatial data to latent space. We chose an autoencoder approach over classical dimensionality reduction strategies like principal component analysis, UMAP or t-SNE due to autoencoders excelling at learning nonlinear and complex representations, which could be essential for optimizing multiple ABM parameters. Other advantages include that new data can be passed directly through the encoder instead of rerunning an algorithm like t-SNE, and the latent space mapping is less sensitive to hyperparameters compared to UMAP. Our current convolutional autoencoder smoothed images in which single lymphocyte cells might be removed within the latent space. This effect could explain for instance the lower correlations in the tumor-lymphocyte neighbors, and how more general structures like tumor growth and shape overpowered lymphocyte-related parameters like IMpkill. Alternative architectures, such as variational autoencoders (mapping spatial data onto a distribution instead of a fixed value) or generative adversarial networks, may better preserve fine-scale distributions (Bond-Taylor *et al*., 2021).

Finally, the observed correspondence between ABM parameters inferred from pathology images and bulk transcriptomic features demonstrates that biologically and molecularly relevant processes can be recovered from spatial imaging data alone. While transcriptomics was used here for a posteriori validation, integrating molecular information directly into the learning or optimization framework could further enhance parameter inference and model interpretability (Retzlaff *et al*., 2023). Such integration would be especially relevant given emerging multimodal tumor modeling approaches, where combining spatial and molecular modalities has shown improved performance compared to single-modality models (Liu et al., 2024; Sadée et al., 2025).

With these rapid developments in data acquisition, we believe that setting up complementary computational frameworks to leverage these data will be key to studying the tumor-immune microenvironment. Our ABM optimization pipeline to inform parameter estimation directly with spatial data is a promising first step in setting up a modular workflow integrating different data modalities for spatial computational models within the field of oncology.

## Acknowledgements

The TCGA images used are data generated by the TCGA Research Network: https://www.cancer.gov/tcga.

