## Supplementary Information for "Informing agent-based models with spatial data using convolutional autoencoders"

Wang B.^ab^

Liao C.^c^

Danen E.H.J.^c^

Neubert E.^c^

Eduati F.*^ab^

*=corresponding author

Author affiliations

^a^Department of Biomedical Engineering, Eindhoven University of Technology, PO Box 513, Eindhoven 5600MB, the Netherlands

^b^Institute for Complex Molecular Systems, Eindhoven University of Technology, 5600 MB Eindhoven, The Netherlands

^c^Leiden Academic Center for Drug Research, Leiden University, Leiden, the Netherlands

Contact information

| ***Supplementary Table: Parameter values for different data sets***  *(*) Optimized parameter, range is given instead of fixed value* | | | | |
| --- | --- | --- | --- | --- |
| **Parameter** | **Description** | **Synthetic** | **Tumoroid** | **TCGA** |
| **oneStepDuration** | Time interval for each simulation step | 12 | 1 | 12 |
| **Height, width** | Height and width (number of cells) within simulation grid | 100 | 100 | 25 |
| **nSteps** | Number of steps per simulation | 150 | 72 | 100 |
| **TUpprol** | Tumor proliferation probability | (*) 0.1-0.5 | 0.15 | (*) 0.1 - 1 |
| **TUpmig** | Tumor migration probability | 0.05 | 0.0005 | 0.05 |
| **TUpdeath** | Tumor death probability | 0.05 | 0.05 | 0.05 |
| **TUpmax** | Maximum number of times tumor can proliferate | 10 | 10 | 10 |
| **TUdamageThresh** | How much damage tumor requires before lymphocyte successfully killed the cell | 2 | 2 | 2 |
| **IMkmax** | Maximum number of times lymphocyte can kill before becoming exhausted | 15 | 15 | 15 |
| **IMpmax** | Maximum number of times lymphocyte can proliferate | 5 | 5 | 5 |
| **IMpmig** | Lymphocyte migration probability | 0.8 | 0.9 | 0.8 |
| **IMpkill** | Lymphocyte killing probability | (*) 0.1 - 0.8 | (*) 0.1 - 0.8 | (*) 0.1 - 1 |
| **IMpprol** | Lymphocyte proliferation probability | 4.6289e-4 | 4.6289e-4 | 4.6289e-4 |
| **IMpdeath** | Lymphocyte death probability | 1.5173e-4 | 0.2 | 1.5173e-4 |
| **IMinfluxProb** | Lymphocyte influx probability | 0.1 | 0.2 | (*) 0.1 - 1 |
| **IMinfluxRate** | Number of lymphocytes that enter simulation during influx | 1 | 1 | 2 |
| **IMrateDynamic** | Scaling parameter to increase number of lymphocytes entering simulation during influx w.r.t. tumor cells | 0.0075 | 0.0075 | 0.0075 |
| **IMrwalk** | Lymphocyte random walk probability to distinguish between directed and random migration | (*) 0 - 1 | 0 | 0.5 |
| **IMinfluxEdge** | Whether lymphocyte influx can only occur at edge of simulation grid (True) or at random throughout entire grid (False) | True | False | False |

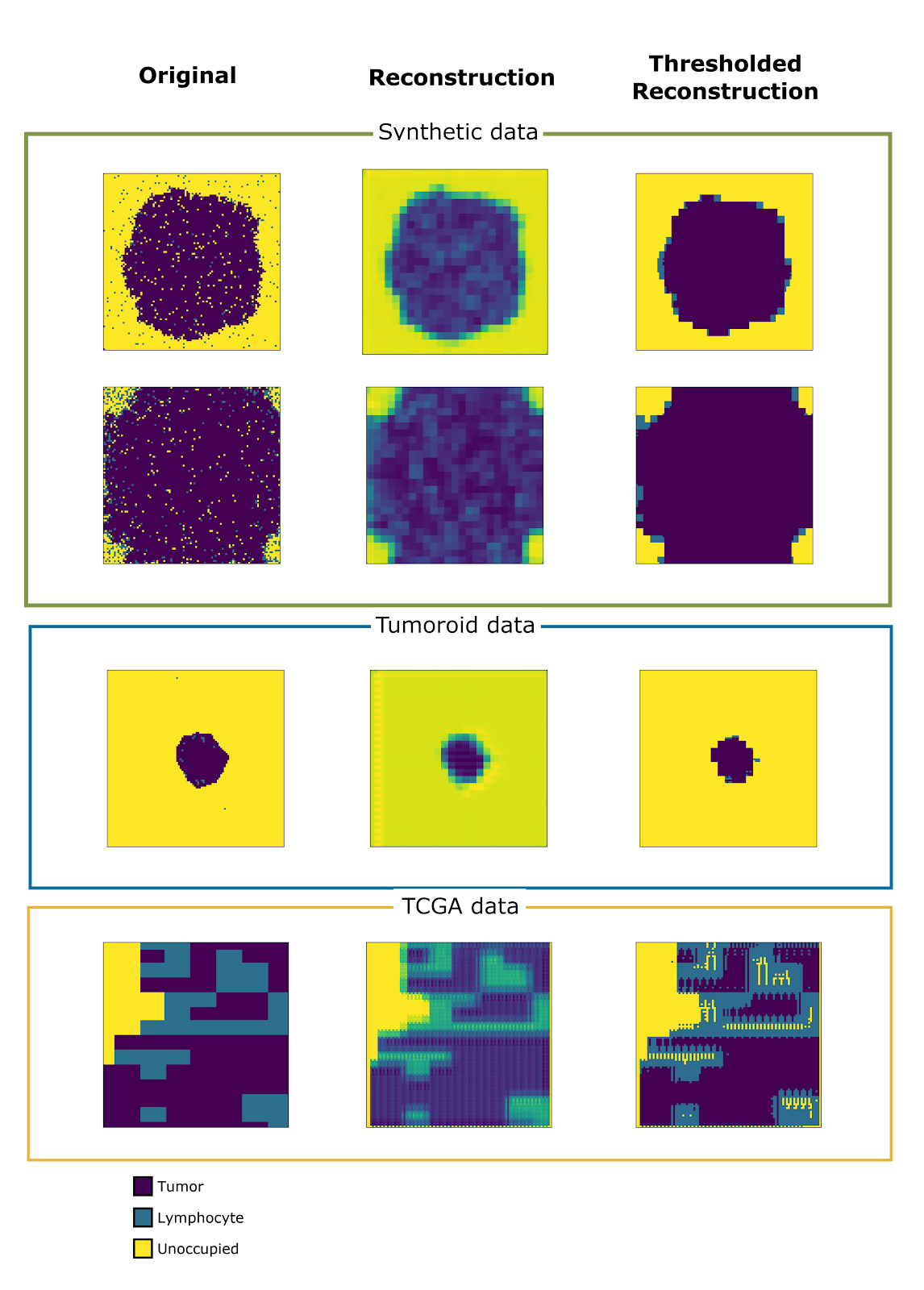

**Supplementary Figure 1: Reconstruction of images with trained autoencoder**. Examples are visualized for different data sets, including original and reconstructed (by trained autoencoder) image and a thresholded reconstruction image. The empirical thresholding categorized the continuous values from the reconstruction into the three values (representing tumor, lymphocyte and unoccupied grids) present in the original images.

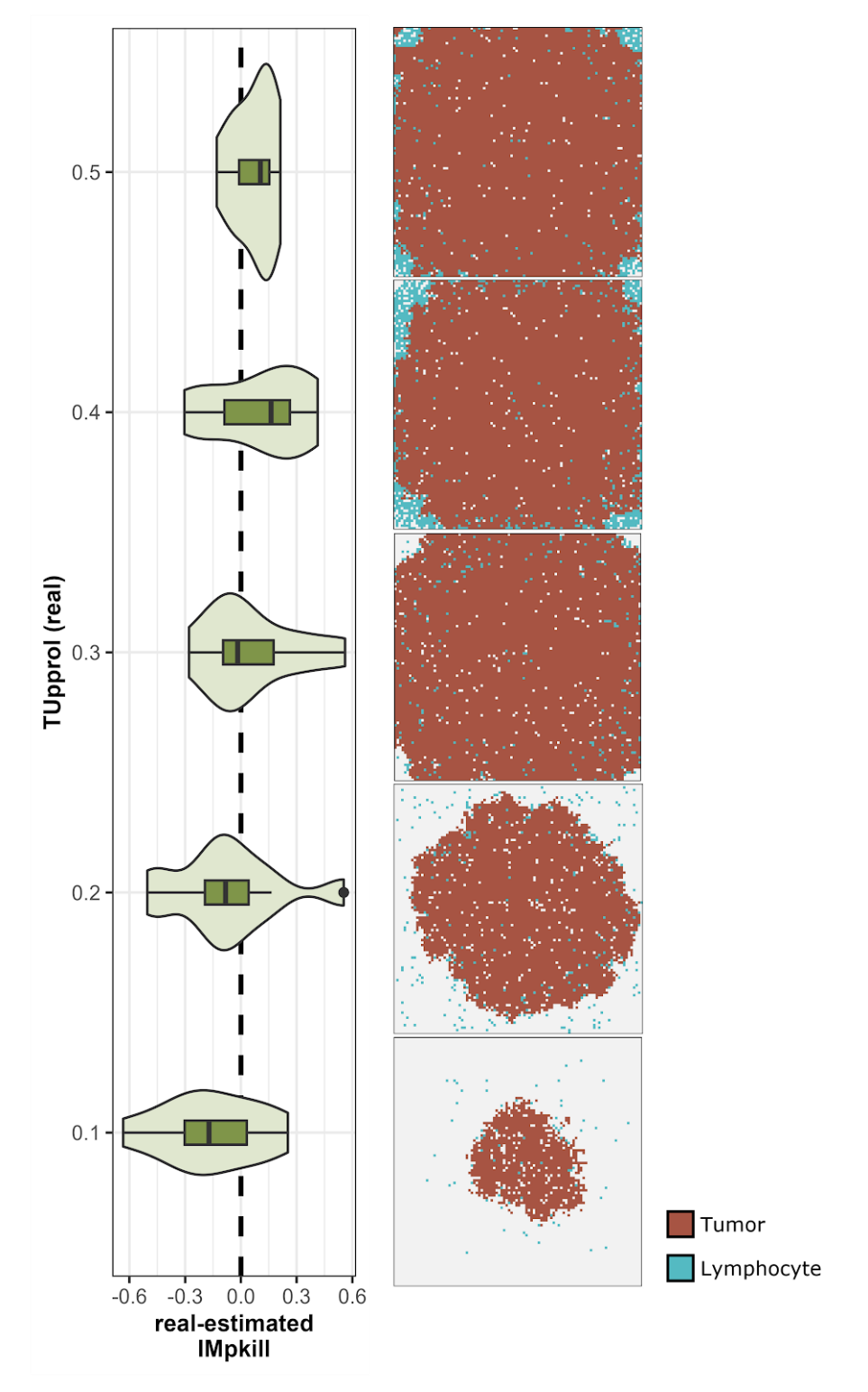

**Supplementary Figure 2:** Effect of TUpprol on differences between ground-truth and estimated IMpkill values. Violinplots represent the difference between ground-truth (real) and estimated IMpkill values, where positive values denote for instance an underestimation of IMpkill. Example spatial outputs are also provided for different TUpprol with IMpkill = 0.4 and IMrwalk = 1.

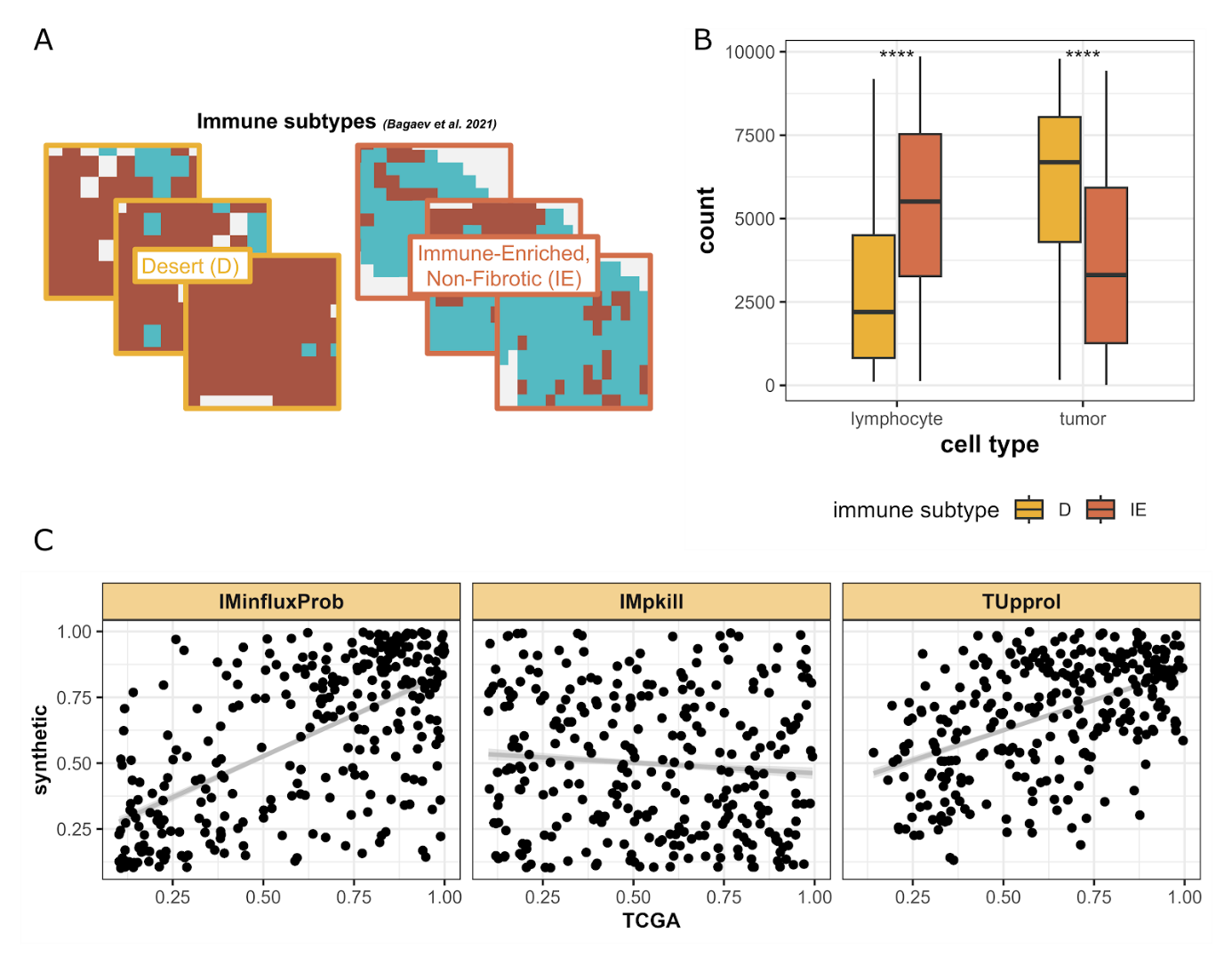

**Supplementary Figure 3: TCGA omics-derived features. (A)** Example patches for each immune subtype (desert, immune-enriched) **(B)** Cell counts of lymphocytes and tumor cells for each immune subtype, with corresponding hypothesis testing for significant differences in means. **(C)** Comparison of estimated IMinfluxProb, IMpkill and TUpprol parameter values with either the autoencoder trained on TCGA data or synthetic data.
